# Multi-scale early warning system for influenza A spillovers

**DOI:** 10.1101/2025.05.24.655955

**Authors:** Tommaso Alfonsi, Anna Bernasconi, Matteo Chiara, Stefano Ceri

**Author notes:** These authors contributed equally to this work (Tommaso Alfonsi, Anna Bernasconi).

## Abstract

Spillovers of Influenza A viruses into farmed animals and humans have the potential to trigger epidemics or even global pandemics. We introduce FluWarning, a highly efficient and elegant computational method based on anomaly detection of codon bias and dinucleotide composition for early flagging of divergent viral HA segments. Applied to H1N1 specimens collected between 2008-2009, FluWarning accurately identifies the emergence of pdm09 - the virus that caused the 2009 flu pandemic; warnings precede the observed progression of the pandemic. Applied to H5N1 specimens collected between 2019 and 2025, FluWarning flagged the emergence of the B3.13 genotype, linked to a spillover event in dairy cows in the United States. In summary, FluWarning is an effective multi-scale warning system for IAVs, detecting spillovers with few available sequences.

**One-Sentence Summary:** Early detection of zoonoses using a few viral sequences is critical for the timely implementation of mitigation strategies.

---

Early detection of changes in the viral genome is paramount for the development of warning systems and the implementation of mitigation strategies. Leaps across different species have been shown to be increasingly frequent (*1*) and can escalate to global health emergencies. Evolutionary breakthroughs may lead to sustainable transmission in a new ecological niche and eventually spillover to further species (as previously observed for Ebola (*2*) or coronaviruses (*3*)).

In information science, conflict and disruption are typically observed through anomaly detection methods; anomalies are carriers of critical information, such as extreme weather conditions, fault tracking, and fraud activities. Several surveys of anomaly detection methods have been proposed previously (*4, 5*), also for temporal data (*6*). Here we introduce FluWarning, a novel computational approach that applies anomaly detection to genomic features, flagging viral specimens showing critical changes with respect to the genome of cognate viral samples. We selected `stray` (*7*), a powerful unsupervised method applied to high-dimensional data.

We use genome sequences retrieved from the Global Initiative on Sharing All Influenza Data (GISAID, (*8*)); we focus on a few fundamental metadata and the hemagglutinin (HA) segment, which is the most commonly sequenced one and modulates host specificity (*9*). However, the method is general and adapts to any collection of IAV sequences available at a surveillance center; the method is also multi-scale, as it can detect individual anomalous sequences.

We previously developed several computational tools to track viral genomic evolution; specifically, we observed single-point mutations across time, on which clustering (*10*), divergence (*11*), or likelihood-based approaches (*12*) allowed us to detect major evolutionary changes (e.g., variants and recombinations) in SARS-CoV-2 genomes. Here we leverage codon usage bias and dinucleotide frequency to capture significant changes in genome composition. Each sequence is encoded in a 75-dimensional feature vector, and the `stray` algorithm is applied to detect sequences with divergent features, which are individually or globally assessed. FluWarning does not aim to predict evolutionary trajectories, as in other works based on anomaly detection (*13, 14*), but is capable of capturing key epidemiological events at their early stage.

FluWarning is configurable by time window and geographic location of interest (from continents to local surveillance units). The method runs in a streamlined, reproducible pipeline (*15*). The results are organised in a user-friendly data mart (*16*) that allows data navigation and selection of specific countries, weeks, and infected host species.

FluWarning has been designed and validated on the paradigmatic case of the 2009 H1N1 pandemic (pdm2009), since progression and all marker events are well-known. The method was then employed in the compelling case of the H5N1 influenza virus, which has recently caused an epidemic in dairy cattle in the United States (*17*) and might be close to reaching pandemic potential according to recent studies (*18*). Our work builds upon a recently published, systematic analysis of H5N1 in North America (*19*).

## Results

### Data collection

From GISAID EpiFlu (*8*), we downloaded 3,387 H1N1 specimens collected from mammalian hosts between July 28^*th*^, 2008 and January 10^*th*^, 2010, and 14,352 H5N1 specimens from any host between January 1^*st*^, 2019 and May 5^*th*^, 2025. For both datasets, we downloaded only specimens collected in North America. After our data curation pipeline (see *Materials and Methods*) we obtained: 3,034 H1N1 sequences (96.4% from human hosts and 3.6% from swine) and 12,568 H5N1 sequences (80.4% from wild birds, 5.7% from domestic birds, 13.5% from non-human mammals, and 0.4% from human hosts).

### Feature Selection

Several studies highlighted how IAVs from different hosts display different genome compositions in terms of di-nucleotides (*20, 21*) and codon usage preferences (*22–26*). The data processing pipeline computes 75 numeric features for every sequence. The first 59 features correspond to the Relative Synonymous Codon Usage (RSCU, (*27*)) of all the codons, excluding the three stop codons (TAA, TAG, TGA), ATG (Met) and TGG (Trp) since the lack of alternative synonymous codons. Further, 16 features describe dinucleotide frequencies of all the permutations *P*(4, 2), with repetitions of the 4 nucleotides, normalized by genome composition.

### Anomaly Detection in High Dimensional Data

`Stray` (Search and TRace Anomaly, (*7*)) focuses on anomaly detection in high-dimensional data; given a set of items, anomalies are defined as items whose distance gap to typical data is significantly larger than the distance between typical items. In our application of `stray`, each item is a vector of 75 variables; items are sorted according to a total order by `CollectionDate` and then by `AccessionID`.

The algorithm is applied to moving windows consisting of *N* items. At each new window *W*_*i*_, we drop the first item of window *W*_*i*–1_ and we add one item *n*_*i*_, denoted as *new item*; we refer to the collection date of the new item as *characteristic date* of the window. Note that the new item in the window *W*_*i*_ is included in the subsequent *N* – 1 windows, where it progressively scales up to becoming the first item, and then is finally removed.

In `stray`, an important, user-defined parameter is *k*; an item is deemed an outlier when the distance to its *k*-th nearest neighbor is above a data-driven threshold, termed “gap”, automatically computed (see *Materials and Methods*); *k* can be interpreted as the maximum size of micro-clusters detected as anomalies. For each window *W*_*i*_, `stray` produces a vector *𝒱*_*i*_ of *N* binary values *v*_*ij*_, where *v*_*ij*_ = 1 indicates that item *j* in the window *i* is anomalous. Finally, a window is associated with a *warning* if the new item is anomalous, i.e. *v*_*i,N*_ = 1.

### Analysis for epidemic/pandemic events

Fig. 1A shows `stray` applied to 51 consecutive windows, representing a complete substitution of blue items (baseline viral population) with red items (new viral population). We consider *N* = 50 and *k* = 10; `stray` is executed 51 times, starting from 50 blue items (blue), progressively replaced by red items. Note that the blue item in position 1 of window 0 is discarded in window 1, making space for an item in position 50 of window 1. This item is an anomaly, as its distance from all other items is greater than a data-defined gap; all anomalies are rendered with a black triangle.

**Figure 1.**
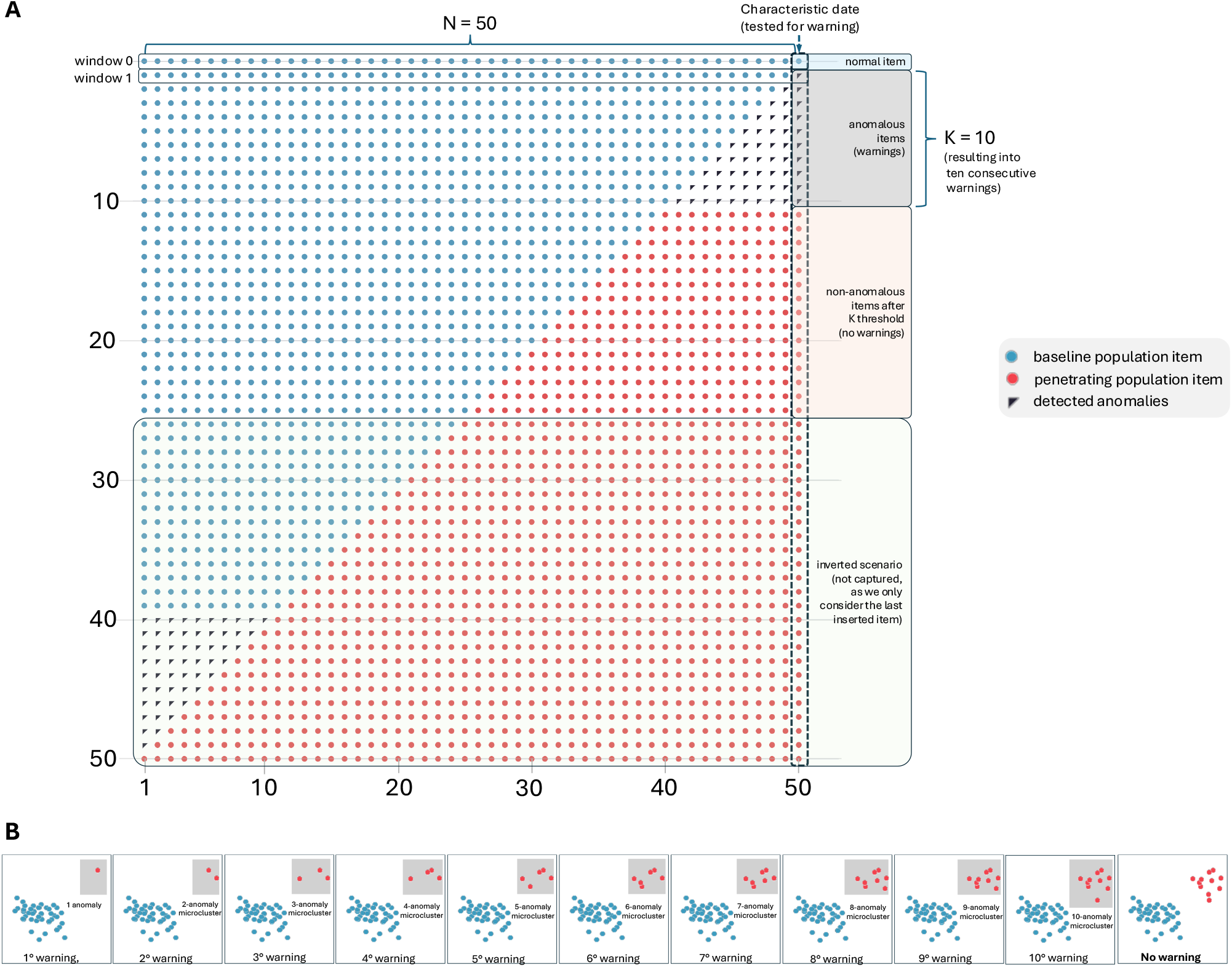
Stray method explained. (**A**) Complete substitution of blue items (a baseline viral population) with red items (a new viral population). With N = 50 and *k* = 10, `stray` is executed 51 times, starting from a window that contains 50 normal items (blue). Anomalies occur only when red items, that progressively form a micro-cluster, are below the user-defined threshold *k* - then, in windows 1 through 10, all are associated with a warning. At window 11, a cluster of size 11 is created, red items cannot be considered anomalous because they have created a sufficiently large cluster. Then, the population of red items grows up to window 40, where blue items are a minority; they create a micro-cluster of size 10, and all blue items become anomalies. However, as we are only interested in new items (in position 50), no warning is thrown. (**B**) shows an “ideal” case where blue and red sequences are well-separated in the 75-dimensional space. Here, we capture distances on two highly-representative dimensions (e.g., the two principal components after dimensional reduction). Red micro-clusters are recognized as anomalous until their size reaches *k* = 10.

Anomalies occur only when red items, that progressively form a micro-cluster, are below the user-defined threshold *k*. Thus, anomalies progressively occur only for windows 1 through 10, all associated with a warning. At window 11, a cluster of size 11 is created, and for each red item, the 10th-neighbor distance drops; red items are no longer anomalies as they belong to a sufficiently large cluster (see Fig. 1B). Then, the substitution of blue items with red items continues, up to window 40. Here, blue items are a minority; they create a micro-cluster of size 10 and become anomalies. However, as we are only interested in new items, no warning is thrown.

#### Sensitivity and specificity analysis

We prepared an *ad hoc* dataset with 1) *N*-sized baseline group (called G1), drawn from an initial set of 130 ordered items (called X1); and 2) *N*-sized penetrating group (called G2), drawn from a different set of 100 ordered items (called X2). The two sets X1 and X2 represent viral sequences collected in the same epidemic year, but from different viral clades, retrieved from GISAID; individual IDs are reported in our Zenodo repository (*15*); each sequence is summarized by 75 extracted features representing its HA viral genome. We build the noise experiments by randomly replacing respectively 10%, 20%, or 30% of the items of G2 with elements of X1 (termed *noise-items*) not used in G1.

Table 1 reports results, with *N* = (50, 100) and *k* = (10, 15); a complete version of the experiments with 15 combinations of *N* and *k* is in Table S1 - but, as discussed in our H1N1 pandemic analysis, smaller *N, k* pairs are not recommended for epidemic/pandemic analyses. Each table line represents a specific combination of *N, k*, and noise level (percentage over G2 and absolute number of *noise-items*). For each combination, we run FluWarning 50 times and average the results, collected in the last four columns, which represent: 1) the number of warnings – identical to *k* in all cases; 2) the window number in which the last warning was detected (denoted as Last Warning Ordinal, LWO); 3) sensitivity, defined as *k*/*LWO* (i.e., the ideal last warning ordinal divided by the actual last warning ordinal – capturing the ratio of true positives over all the warnings); and 4) specificity, defined as (*N* + 1 – *LWO*)/(*N* + 1 – *k*) (i.e., the ideal ordinal from which non-warning items are detected, divided by the actual non-warning ordinal – capturing the ratio of true negatives over all the warnings).

**Table 1.** Sensitivity and specificity analysis, with *N* = (50, 100) and *k* = (10, 15). Each line represents a specific combination of *N* (column 1), *k* (column 2), and noise level as a percentage over G2 (column 3). The absolute number of *noise-items* is also reported (column 4). For each combination, we run FluWarning 50 times; we report the number of warnings (column 5); averages of the window number in which the last warning was detected, denoted as Last Warning Ordinal - LWO (column 6); sensitivity (column 7); and specificity (column 8).

| N | K | %Noise | #NoiseItems | #Warnings | LWO | Sensitivity | Specificity |
| --- | --- | --- | --- | --- | --- | --- | --- |
| 50 | 10 | 0 % | 0 | 10 | 10.00 | 1 | 1 |
|  |  | 10 % | 5 | 10 | 11.16 | 0.901 | 0.972 |
|  |  | 20 % | 10 | 10 | 12.68 | 0.798 | 0.935 |
|  |  | 30 % | 15 | 10 | 13.84 | 0.736 | 0.906 |
|  | 15 | 0 % | 0 | 15 | 15.00 | 1 | 1 |
|  |  | 10 % | 5 | 15 | 16.70 | 0.900 | 0.953 |
|  |  | 20 % | 10 | 15 | 18.60 | 0.812 | 0.900 |
|  |  | 30 % | 15 | 15 | 21.10 | 0.719 | 0.831 |
| 100 | 10 | 0 % | 0 | 10 | 10.00 | 1 | 1 |
|  |  | 10 % | 10 | 10 | 11.00 | 0.915 | 0.989 |
|  |  | 20 % | 20 | 10 | 11.98 | 0.849 | 0.978 |
|  |  | 30 % | 30 | 10 | 14.52 | 0.709 | 0.950 |
|  | 15 | 0 % | 0 | 15 | 15.00 | 1 | 1 |
|  |  | 10 % | 10 | 15 | 16.52 | 0.913 | 0.982 |
|  |  | 20 % | 20 | 15 | 18.22 | 0.829 | 0.963 |
|  |  | 30 % | 30 | 15 | 21.70 | 0.706 | 0.922 |

As expected, LWO increases with contamination, while sensitivity and specificity decrease from the perfect values 1 (no contamination); however, their values are acceptable also with 30% contamination, showing that the method is robust. Note that, in any warning system, a few false positives are not a concern, as all sequences that triggered a warning must be further explored.

#### Data mart-based monitoring

FluWarning is distributed as software supporting (1) *data processing* and (2) *data exploration*. In (1), the software receives a stream of sequences (e.g, downloaded from GISAID by an authorized user or provided from a surveillance laboratory). After sequence alignment, feature selection, and anomaly detection, a database including warning-associated sequences and their metadata is collected within a (SQLite) database. In (2), the database feeds a *data mart*, explorable through a web-based application (Streamlit), packaged in an easy-to-install container (Docker), see (*15*). The data mart supports the inspection of aggregate counts of warnings across selected hosts, locations and periods, as well as the extraction of single sequences metadata; the software is developed to facilitate installation and use.^1^

### The H1N1 pandemic (2008-2009)

We used the 2009 H1N1 pandemic to identify the most appropriate settings for FluWarning for the detection of epidemiologically important events. The pdm09 virus, which caused the pandemic, originated in late 2008 in Mexico, but was first isolated and sequenced on April 14th, 2009, by the Centers for Disease Control and Prevention (*28*). Following a rapid increase in flu cases associated with pdm09, the World Health Organization (WHO) declared a “public health emergency of international concern” on April 25th, 2009. By June, pdm09 completely replaced other seasonal H1N1 flu strains, and the outbreak was declared a pandemic (*29*).

We analyzed 3,034 H1N1 genome sequences obtained from human hosts and collected between August 2008 and January 2010. Data were aggregated in non-overlapping intervals of two weeks. Each interval has a *Starting Week Number W*_*i*_ (with incremental numbers, per year), a corresponding *Start Date D*_*i*_, and includes all the sequences with a collection date included in the interval [*D*_*i*_, *D*_*i*+1_). To identify the most appropriate settings for the identification of the emergence of a novel virus in FluWarning we visually inspected the total number of warnings emitted by the system at every interval *W*_*i*_, according to different combinations of *N* and *k*. As shown in Fig. S1, with *N* = (50, 100) and *k* = (10, 15) a sharp peak of warnings was observed, specifically at intervals ranging from 2009-11 to 2009-20. These intervals span from March 12th, 2009 to May 13th, 2009 and align exactly with the initial emergence and spread of pdm09. Based on these observations, only values of *N* = (50, 100) and *k* = (10, 15) were considered for epidemic/pandemic scale analyses.

A Fisher exact test (see *Materials and Methods*) was applied to identify *W*_*i*_ associated with a statistically significant increase in the number of detected anomalies; p-values were corrected with the Benjamini-Hochberg procedure to control the false discovery rate. We reasoned that intervals associated with a statistically significant increase (corrected p-value ≤ 0.05) in the number of detected anomalies should reflect key epidemiological events as -for example-the spread of a new virus. From now on, we will refer to these intervals as *Super Warning*s.

Results are shown in Fig. 2A–C. For all four configurations of `stray`, *Super Warning*s (see dark blue cells) were emitted from 2009-13 to 2009-21. The first pdm09 sequence in the dataset was collected on March 12th, 2009 (week 2009-11) and was identified as a warning by `stray` in all four configurations. Four *Super Warning*s were instead emitted at 2009-13; by this interval of time, the number of pdm09 sequences in the GISAID database increased to 14, collected in Mexico (n=7) and California (n=7), consistent with the epidemiological history of the H1N1 pandemic of 2009.

**Figure 2.**
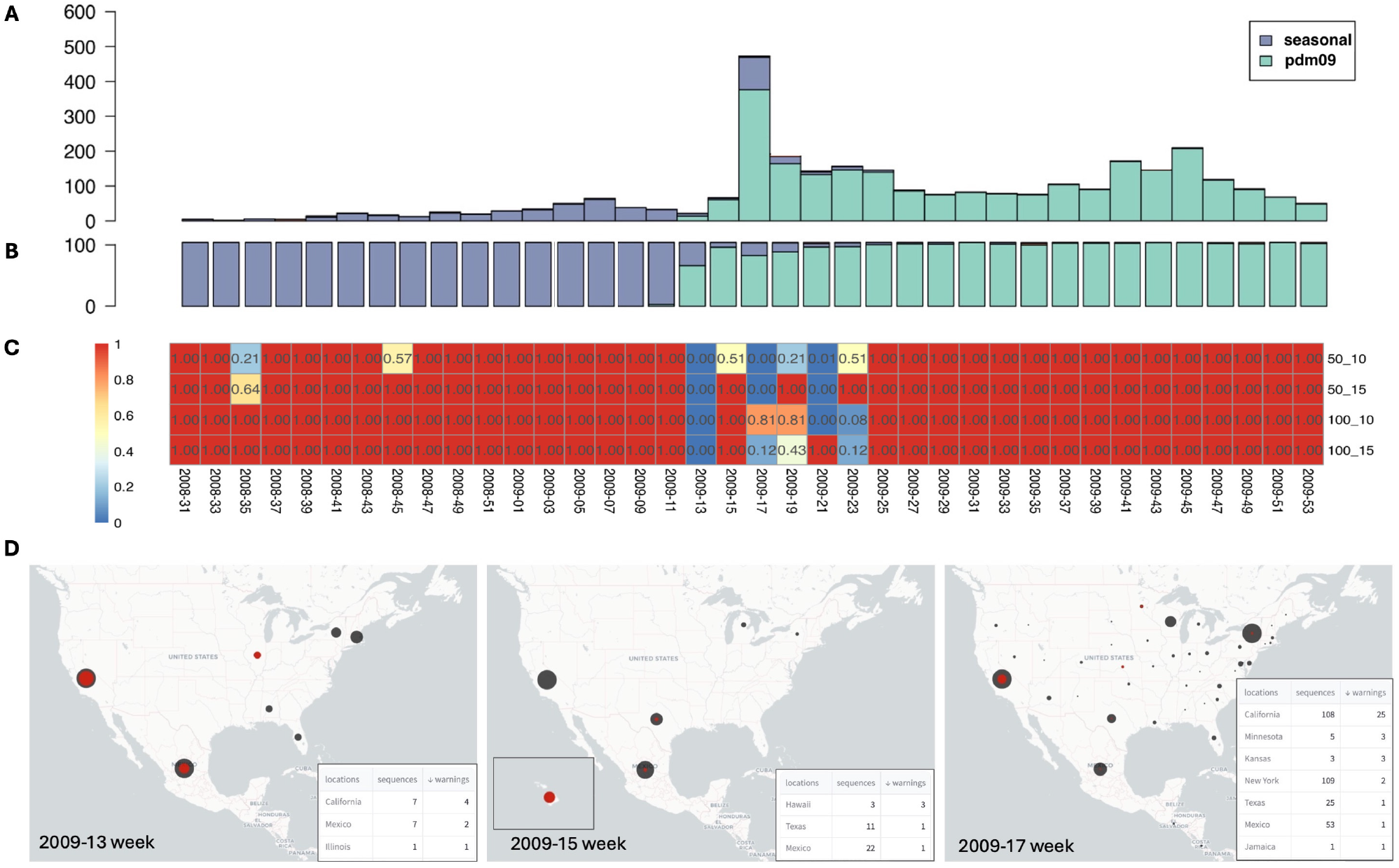
Circulation of H1N1 strains from 2008-33 to 2009-53 and warnings by stray. (**A**) Barplot showing the total number of H1N1 sequences classified as “seasonal-flu” or “pdm09”, according to the GISAID EpiFlu database, at every time interval. (**B**) Relative proportions (percentage with respect to the total number of sequences) of seasonal and pdm09 sequences. (**C**) Heatmap (with `stray` settings *N* = 50, 100 and *k* = 10, 15) displaying the FDR corrected p-values for the statistically significant increase in the number of warnings. Time intervals are shown on columns, parameters of `stray` on the rows. (**D**) Maps drawn for North and Central America, representing warnings during the 2009-13 to 2009-17 weeks period. One can note the diffusion, initially in Mexico and Southern California (as it can be read in location metadata), up to other States of the US.

Interestingly, starting from week 2009-15 – when pdm09 established itself as the most prevalent H1N1 clade in the dataset – the majority of the warnings emitted by the system flagged seasonal flu viral isolates. By week 2009-25, pdm09 accounted for (almost) the totality of H1N1 samples, and the number of warnings identified by `stray` at any subsequent intervals dropped to levels comparable to those observed before the emergence of pdm09 (see Fig. S2 and Data S1).

#### Data mart-based monitoring

To showcase the power of our data mart, consider Fig. 2D, obtained by setting *N* = 50 and *k* = 10, which illustrates the locations of warnings. In bi-week 2009-13/14 (March 23rd - April 5th, 2009), warnings occurred in California (4 warnings out of 7 isolates), Mexico (2/7), and Illinois (1/1); in bi-week 2009-15/16 (April 6th-19th, 2009), warnings occurred in Hawaii (3/3), Texas (1/11), and Mexico (1/22). By week 2009-17/18 (April 20th - May 3rd, 2009), a significant number of warnings occurred in California (25/108), and US-collected warnings spread to Minnesota, Kansas, New York, and Texas, marking the start of the pdm09 pandemic. Collectively, these results indicate that FluWarning correctly recapitulates the main events of the 2009 H1N1 pandemic and flags the emergence of new viral clades/strains almost in real time. These considerations prompted us to apply the same analytical workflow to H5N1.

### The H5N1 2.3.4.4b clade (2020-2025)

Highly pathogenic (HPAI) strains of H5N1 IAVs are considered a constant pandemic threat since their numerous outbreaks in mammalian species, including sporadic cases of human infection with reported high mortality rates (*30*). Originally described and isolated in 2013-2014 in Asia, the HPAI H5N1 2.3.4.4b clade recently crossed continental barriers and became increasingly frequent in Europe, North America, and South America (*31*). Besides infecting wild and domestic birds, 2.3.4.4b was associated with several distinct spillovers to mammals throughout 2020 to 2024 and recently caused a large epidemic of avian flu in dairy cattle in the United States, causing significant concern for human health (*17*). A single genotype, labelled B3.13, was associated with the 2024 H5N1 dairy cattle flu epidemic; the first isolate of B3.13 was collected from a dairy cow in Texas on 2024-03-13 (*19*).

We considered a total of 12,568 HA segment sequences and associated metadata of H5N1 collected between September 15th, 2020 and March 3rd, 2025 in North America^2^. A total of 1,035 distinct warnings were emitted by FluWarning in at least one of the *N* = 50, 100 and *k* = 10, 15 configurations (see Fig. S3 and Data S1); sequences assigned to the B3.13 genotype accounted for 55.36% of the total number of warnings and 84.8% of the warnings emitted by FluWarning between 2024-11 and 2024-29. These windows were associated with *Super Warning*s according to all four selected configurations of `stray`, see Fig. 3C (complete heatmap in Fig. S4). As observed from the breakdown of the circulation of 2.3.4.4b genotypes in North America, displayed in Fig. 3A-B, this interval aligns almost exactly with the emergence and spread of B3.13. Several other genotypes, including for example B2.1, B3.2, B3.6, display a widespread and sustained circulation in North America (estimated prevalence ≥ 0.2 for more than 4 weeks), but none of these highly frequent genotypes exhibited a statistically significant association with *Super Warnings*. Instead, a marked increase in the number of warnings was clearly associated with B3.13, as observed in Fig. S3.

**Figure 3.**
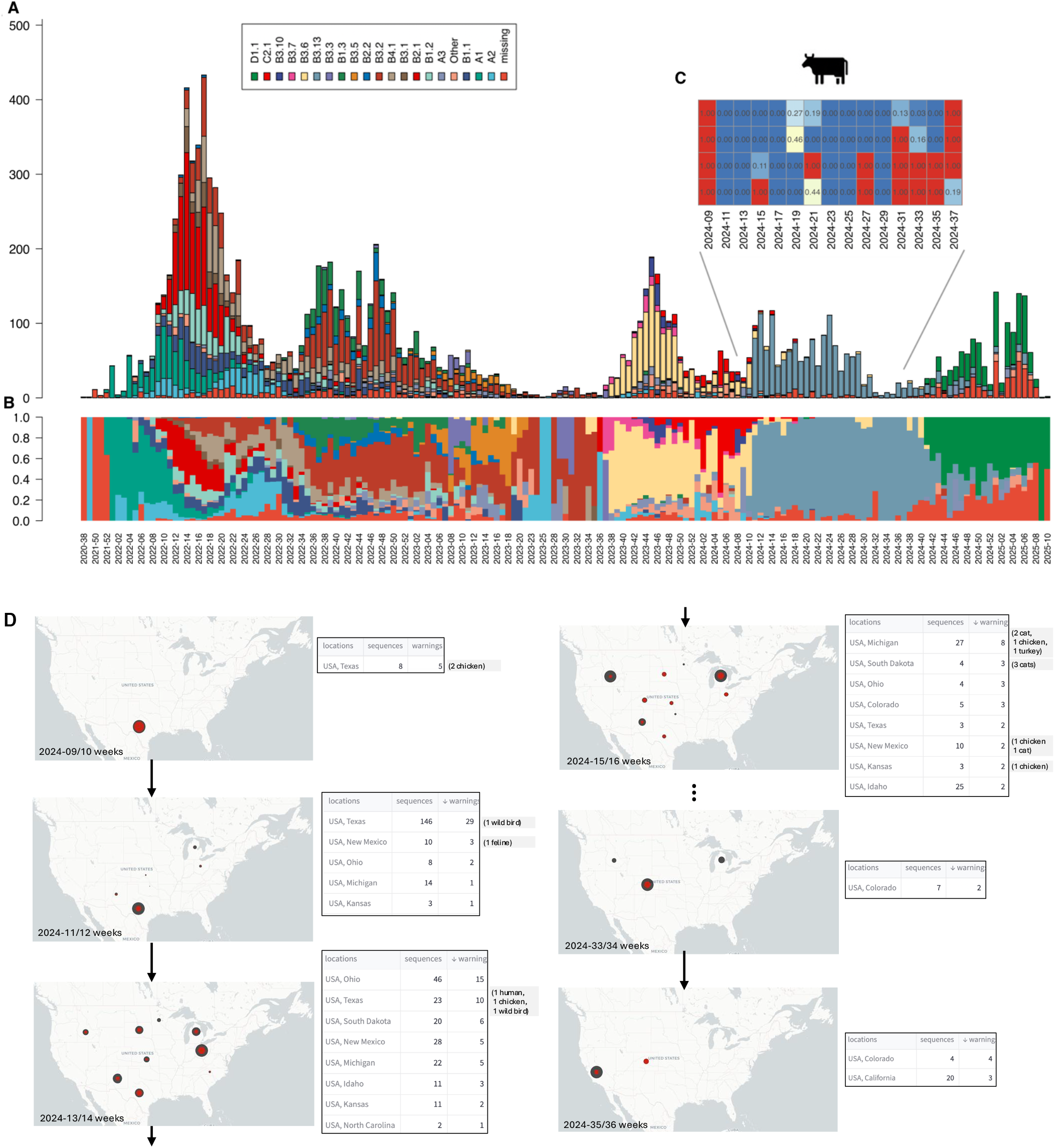
Circulation of 2.3.4.4b H5N1 strains from 2020 to 2024 and warnings by stray. (**A**) Barplot showing the total number of H5N1 sequences at every time interval, classified according to the genotypes recently associated with H5N1 in North America. (**B**) Relative proportions (percentage with respect to the total number of sequences) of genotypes, shown to facilitate the direct comparison of their prevalence. (**C**) Heatmap displaying the FDR-corrected p-values for the statistically significant increase in the number of warnings. Only the intervals associated with the diffusion in non-human mammals (dairy cattle) is displayed (complete analyses in Fig. S4). Time intervals are shown on columns. Parameters of `stray` on the rows. (**D**) Maps drawn for North and Central America, representing warnings during the 2024-09 to 2024-36 weeks period. One can note the diffusion, initially in Texas (week 9-10), then also in New Mexico, Ohio, Kansas and Michigan (week 11-12), then also in Idaho, North Carolina and South Dakota (week 13-14), then also in Colorado (week 15-16); we move then to weeks 33-36 to show the offspring of warnings in California, first presented in weeks 35-36.

#### Data mart-based analysis for individual B3.13 sequences

FluWarning can be applied to surveillance on smaller scales (US States in our dataset, but -in principle-regional surveillance centers); with more granular analysis, `stray` can be exploited with lower *k* parameters. When focusing only on B3.13 sequences, a progressive movement of warnings (using *N*=50 and *k*=1) is observed in the same bi-weekly intervals – see Fig. 3D, showing subsequent US maps and the corresponding B3.13 warning counts (automatically generated by our data mart). Each panel shows geo-located warnings (red circle) included in collected sequences (black circle). Numbers are explicitly indicated in the tables below; grey cards specify warnings thrown by sequences found on hosts other than dairy cows.

Starting March 7th, 2024, Texas collected two warning sequences from chicken, then on March 10th, 2024, three from dairy cows; the following intervals see an increase in warnings in Texas (2 wild birds, 1 chicken, and 1 human in 2024-13 ID: “*EPI_ISL_19027114_A/Texas/37/2024*”). At weeks 2024-11/12, we observe warnings in New Mexico (March 15/16th), Ohio (March 15th), and subsequently Kansas (March 20th), New Mexico (feline on March 20th), and Michigan (March 21st). In the interval 2024-13/14, warnings were also recorded in Idaho (March 27th), North Carolina (April 4th), and South Dakota (April 5th). In the interval 2024-15/16, warnings appear in Colorado (April 10th). The last two maps depict a much later period, showing the emergence of warnings in California (2024-35/36), precisely on August 26th, 2024 (ID: “*EPI_ISL_19387789_A/dairy_cow/USA/24_024712-003/2024*”).

Warnings emitted for 5 US States are shown in Fig. S5. For every row, we report bi-weekly counts, according to all possible assignments of parameters *N* and *k*. Note the peculiar situation of California: warnings are emitted for a long period, from August 2024 to February 2025, and then no sequence appears as collected, in spite of the reported critical situation of dairy cattle (*19*).

#### Cluster and mutation analysis

The HaploCoV (*32*) software was applied to delineate clusters of similar HA sequences in 2.3.4.4b. A total of 10 distinct groups were formed (Fig. 4A). Interestingly, 3 of these 10 groups (HA.N1, HA.N2, and HA.N10) were composed almost exclusively of sequences assigned to the B3.13 genotype, while the B3.6 genotype -ancestor of B3.13 according to (*19*)-corresponded with the HaploCoV cluster labelled HA.N3.

**Figure 4.**
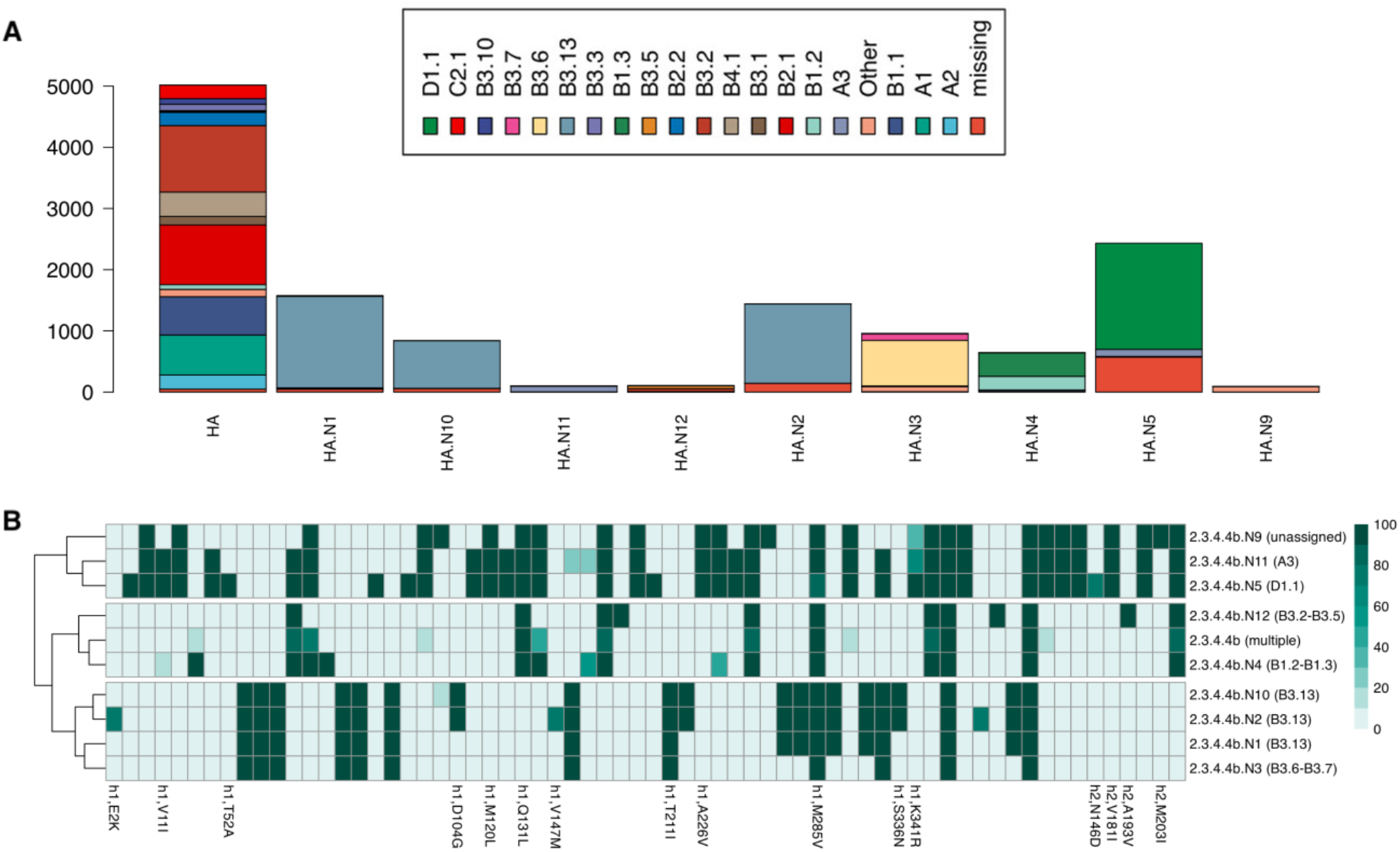
Number of sequences of each genotype corresponding to a HaploCoV-defined cluster and characteristic mutations of HaploCoV clusters. (**A**) For every HaploCoV cluster, the barplot displays the total number of sequences from each distinct genotype assigned to that cluster. Clusters HA.N1, HA.N10, and HA.N2 are formed almost exclusively by sequences assigned to the B3.13 genotype. Similarly, a clear correspondence between HA.N3 and B3.6 can be observed. (**B**) The heatmap illustrates the prevalences (% min 0 max 100) of characteristic nucleotide variants in each distinct HaploCoV cluster. HaploCoV clusters are reported on the columns; nucleotide variants on the rows. For nucleotide variants associated with non-synonymous substitutions, the predicted amino acid substitution is reported (complete annotation in Data S1). Consistent with phylogenetic analyses, clusters associated with B3.6 (HA.N3) and B3.13 (HA.N1, HA.N10, and HA.N2) form a neat and well-separated group and are defined by a similar pattern of characteristic nucleotide variants.

As observed in Fig. S6, the B3.6 group, hosted by wild/domestic birds, was dispersed across many states; HA.N1 was associated prevalently with specimens collected from dairy cattle in Texas, Colorado, Idaho, Michigan, and Minnesota; instead, the HA.N10 and HA.N2 clusters, detected starting from week 2024-35, were observed only in California.

We designate as defining nucleotide variants those observed in at least 50% of the sequences of each HaploCoV cluster; these nucleotide variants were inspected (see Fig. 4B). Consistent with phylogenetic analyses, HA.N3 (B3.6) and HA.N1, HA.N10, HA.N2 (B3.13) shared a similar pattern of defining nucleotide variants and formed a well-defined group. HA.N1, HA.N10, and HA.N2 displayed five additional characteristic nucleotide variants compared to HA.N3; none of them resulted in an amino acid substitution. However, HA.N2 and HA.N10 displayed two additional non-synonymous substitutions S336N and D104G, compared to HA.N1. S336N was previously reported as a virulence marker, with an increased prevalence in viruses isolated from dairy cows (*19*). D104G was recently associated with a marked increase in immune escape (*33*). Taken altogether, these mutations point to preliminary evidence of host adaptation, and according to our analyses, they are prevalently observed in viral isolates collected in California.

## Discussion

Our FluWarning method issues warnings whenever a new viral sequence is significantly different from a baseline of *N* previously collected background sequences; differences are measured in a dimensional space of 75 variables, describing codon preferences and dinucleotide frequencies. The method does not differentiate among warnings; these may reflect spillovers, new clades, or new lineages. Investigation responsibility is passed to users who interpret the warning. We illustrated the robustness of the method by computing sensitivity and specificity with increasing amounts of contamination, starting from an ideal case of replacement of the baseline population.

FluWarning is conceived for supporting surveillance systems that sample domestic and wild animals globally; it is most likely to succeed when applied to pathogens with an existing surveillance network (now in place only for seasonal influenza). Even under the best surveillance systems (like the WHO’s seasonal influenza GISRS system), sequences routinely get backfilled for earlier dates after the sample collection. Therefore, should public health officials run FluWarning over public data, the input would be relatively incomplete and noisy.

Thanks to its light-weight code, user-defined parametrization, and easy-to-inspect data mart analytics, FluWarning has the potential of being used by many laboratories or regional-scale genomic surveillance institutions. The accuracy of our tool shows that broader and deeper support for global surveillance could pay off, enabling significant small-scale discoveries.

## Acknowledgments

We gratefully acknowledge all data contributors, i.e. the Authors and their Originating Laboratories responsible for obtaining the specimens, and their Submitting Laboratories that generated the genetic sequence and metadata and shared via the GISAID Initiative the data on which part of this research is based. The authors are grateful to Ilaria Capua (Johns Hopkins University), the group of Alice Fusaro and Isabella Monne (Istituto Zooprofilattico Sperimentale delle Venezie), and Manuela Sironi (IRCCS Eugenio Medea), for the fruitful discussions inspiring this research.

## Author contributions

TA and AB are co-primary authors; TA performed data collection and processing; TA, AB, and SC adapted `stray` for the purposes of this paper and performed the specificity and sensitivity analysis; TA and AB designed the data mart. All authors discussed the bi-weekly setting and the presentation of H1N1 and H5N1 cases. MC performed the bioinformatic analyses of H1N1 and H5N1, in particular leading to the identification of relevant mutations for H5N1 and devised the statistical test for the identification of *SuperWarning*s. All authors contributed to the writing; AB is the contact author; SC coordinated the research.

## Data and materials availability

Original sequences and metadata are publicly accessible through the GISAID platform (H1N1: EPI_SET_250523as (*34*) and H5N1: EPI_SET_250523tm (*35*)). The list of sample accessions and FluWarning code are provided on our Zenodo repository (*15*).

## Supplementary materials

Materials and Methods

Figs. S1 to S6

Table S1

Data S1

## Footnotes

1 **Note for reviewers**: We provide passwrord-protected access to the data mart storing GISAID viral sequences (see submission letter / Supplementary Materials PDF).

1 We extracted GISAID data until May 5, 2025; 2,895 sequences submitted in 2025 had incomplete geographic and collection date metadata, and could not be evaluated.

