## Supplementary material for "Multi-scale early warning system for influenza A spillovers": Supplemantary Material

**This PDF file includes:**

Materials and Methods

Figures S1 to S6

Table S1

### Materials and Methods

#### Data collection and genome quality filtering

We download from GISAID EpiFlu (*8*) two sets of sequences and their metadata: (1) the H1N1 sequences deposited in North America from mammalian hosts between the July $28^{th}$, 2008 and January $10^{th}$, 2010; and (2) the H5N1 sequences deposited in North America from all available hosts between January $1^{st}$, 2019 and May $5^{th}$, 2025 (however, no North American sequences with complete collection date metadata were deposited after March $3^{rd}$). For each isolate, we require the HA segment to be available and complete, and the date of collection to be precisely specified. The two sets count 3,387 and 14,352 sequences, respectively. Our data cleaning pipeline removes the sequences with a percentage of unknown nucleotide bases > 2%, duplicate `Isolate_ID` and `Isolate_Name`, incomplete `Collection_Date`, or whose length is > ±7% dissimilar from the mode. Then, we extract the coding sequence (CDS) of the hemagglutinin protein, and assign each codon with the corresponding amino acid according to the genetic code; we remove those sequences with incomplete or truncated CDS (i.e., when the length is not a multiple of 3). After this pipeline, we obtain 3,034 H1N1 sequences and 12,568 H5N1 sequences. On average, the CDS studied in the subsequent analyses is 1,701 nb long for H1N1 and 1,703 nb long for H5N1. The distribution of host species shows that 96.4% of the H1N1 sequences are from human hosts and the remaining 3.6% (146 sequences) from swine. For H5N1 sequences, 80.4% of the sequences are collected from wild birds, 5.7% from domestic birds, 13.5% from non-human mammals, and 0.4% from human hosts.

#### Data processing

The data processing pipeline extracts an array of 75 numeric features from the CDS of each input sequence. The first set of features is the logarithm of the Relative Synonymous Codon Usage (RSCU) (*27*), which is computed for all codons except 3 stop codons (TAA, TAG, TGA) and 2 non-synonymous codons (ATG, TGG). This leads to 59 log-RSCU values for each CDS, computed as in Equation S1, where $X_{ij}$ is the frequency of the codon $j^{th}$ encoding the $i^{th}$ amino acid, and

$n_i$ is the number of synonymous codons encoding the $i^{th}$ amino acid. The RSCU allows us to measure the over/under usage of a codon within a genomic sequence, assuming even codon usage between synonymous codons as baseline value. One common pitfall of the RSCU is that the values representing under-usage are shrunk between 0 and 1, while those representing over-can reach values well beyond 2 or 3. To balance the range of values for expressing the under/over-usage, the RSCU is transformed through the logarithm function to base $n_i$.

$$log\text{-}RSCU(j) = log_{n_i}\left(\frac{X_{ij}}{\frac{1}{n_i}\sum_{j=1}^{n_i} X_{ij}}\right) \quad \text{(S1)}$$

The second set of features derives from the dinucleotide frequencies of all the permutations $P(4, 2)$ with repetitions of the 4 nucleotides in each CDS sequence. We report the formula in Equation S2 for the permutation $C, A$ in a CDS of length $N$; a $log_2$ transformation is applied to homogenize the values with those of log-RSCU.

$$log\text{-}dinucleotide(C, A) = log_2\left(\frac{\frac{count(C,A)}{N-1}}{\frac{count(C)*count(A)}{N^2}}\right) \quad \text{(S2)}$$

**More details on `stray`**

The `stray` method is an improved variant of the HDoutlier method (*36*). Essentially, while HDoutlier uses neighbor distance to identify outliers, `stray` uses the $k$-neighbour distance, thus allowing micro-clusters of outliers of size $k - 1$. In addition, it provides a data-driven threshold for anomaly detection. Some aspects of the `stray` method require further explanation. First, as variables with large variance can have a disproportional influence on Euclidean distance calculation (*36*), the input features for `stray` are normalized between [0,1] through a standard min-max scaler right before running the outlier detection algorithm.

The anomalous threshold calculation is an application of Weissman's spacing theorem (*37*) that applies to data distributions covered by the maximum domain of attraction of a Gumbel distribution; note that this requirement is satisfied by a wide range of distributions, including the exponential, gamma, normal and log-normal distributions with exponentially decaying tails (*7*). For the threshold calculation, `stray` adopts the 'bottom-up searching algorithm' defined in (*38*); the threshold calculation is performed under the assumption that the distribution of $k$-nearest neighbors with the maximum gap is in the maximum domain of attraction of the Gumbel distribution.

For simplicity, we consider as output a binary classification; the `stray` algorithm may also assign an anomalous score to each data instance.

The method considers totally ordered data streams for optimizing the coding of the `stray` algorithm (*39*). Our dataset is totally ordered by ⟨depositionTime,isolateID⟩, fitting the `stray` input requirements.

#### Identification of SuperWarnings

Intervals of time associated with a significant increase in the number of warnings were identified by a one-sided Fisher's exact test. Briefly, for every interval, a $2 \times 2$ contingency table was built by placing, on row 1, counts regarding the observed interval, and on row 2, counts regarding all other intervals. Counts in column 1 represent the total number of sequences labeled as warnings, and counts in column 2 represent the total number of sequences not labeled as warnings. The test statistics and the p-value were computed by means of the `fisher.test()` function as implemented by the `stats` package in R. Correction for multiple testing was computed by the `p.adjust()` function from the same software library.

#### Clusters of HA sequences and characteristic mutations

The HaploCoV software (*32*) was used to identify groups (clusters) of sequences that share common patterns of nucleotide variants in the HA segment of H5N1. HaploCoV detects clusters consisting of at least a user-specified minimum number of sequences ($S$), each containing at least $N$ high-frequency nucleotide variants (with a frequency of occurrence in the dataset $\geq$ 1%) not shared with any other group. In this analysis, the parameters were set as $S = 100$ and $N = 3$.

Each cluster is labeled with the prefix .N, followed by a sequential number. Following the approach in (*19*), the PQ705761 sequence (GenBank accession) was used as the reference sequence for the identification of nucleotide variants. For every cluster, characteristic mutations were defined as those observed in at least 50% of the sequences assigned to a clade/subclade.

**Data mart implementation**

Fifteen configurations of `stray`, obtained by setting $N$ and $k$ parameters, are run on the full dataset extracted as described in the 'Data collection and genome quality filtering' section. For each configuration, FluWarning has been run on $N$-sized windows, progressively acquiring a new sequence and dropping a previous sequence, following a complete order built by `CollectionDate` and then `AccessionId`. The outputs of each run (each resulting in either 0 or 1 warning) are aggregated in macro-windows of two weeks, by summing the number of runs that output one warning (see *Results*, H1N1). Bi-weekly counts of warnings make results more readable and applicable to practical, real-world scenarios. These aggregations are stored, generating a large data mart (*16*). While the method is run on the whole dataset, results can be projected on subsets of sequences corresponding, e.g., to specific host types, geographical locations, and time intervals (e.g., sequences collected in `Canada` or `Massachusetts` for `Swine` hosts in the interval `March 2009 - Sept 2009`). Counts are evaluated as absolute values or as percentages of warnings in the observed bi-weekly period. The selected data structure allows us to quickly navigate the number of runs that have been generated by FluWarning. Most importantly, it allows us to keep an up-to-date visual representation (e.g., on geographical maps) of how warnings are distributed worldwide and how they evolve along bi-weekly intervals.

| N | K | %Noise | #NoiseItems | #Warnings | LWO | Sensitivity | Specificity |
|---|---|---|---|---|---|---|---|
| 5 | 1 | 0% | 0 | 2.0 | 2.00 | 0.500 | 0.800 |
| | | 20% | 1 | 2.0 | 2.50 | 0.417 | 0.700 |
| 10 | 1 | 0% | 0 | 1.0 | 1.00 | 1.000 | 1.000 |
| | | 10% | 1 | 1.2 | 1.60 | 0.850 | 0.940 |
| | | 20% | 2 | 1.4 | 2.20 | 0.672 | 0.880 |
| | | 30% | 3 | 1.9 | 3.50 | 0.471 | 0.750 |
| | 3 | 0% | 0 | 3.0 | 3.00 | 1.000 | 1.000 |
| | | 10% | 1 | 3.0 | 3.20 | 0.950 | 0.975 |
| | | 20% | 2 | 3.0 | 3.57 | 0.862 | 0.929 |
| | | 30% | 3 | 3.4 | 4.87 | 0.659 | 0.767 |
| 50 | 1 | 0% | 0 | 1.0 | 1.00 | 1.000 | 1.000 |
| | | 10% | 5 | 1.0 | 1.02 | 0.990 | 1.000 |
| | | 20% | 10 | 1.0 | 1.18 | 0.917 | 0.996 |
| | | 30% | 15 | 1.0 | 1.36 | 0.840 | 0.993 |
| | 3 | 0% | 0 | 3.0 | 3.00 | 1.000 | 1.000 |
| | | 10% | 5 | 3.0 | 3.28 | 0.930 | 0.994 |
| | | 20% | 10 | 3.0 | 3.84 | 0.826 | 0.983 |
| | | 30% | 15 | 3.0 | 4.00 | 0.802 | 0.979 |
| | 5 | 0% | 0 | 5.0 | 5.00 | 1.000 | 1.000 |
| | | 10% | 5 | 5.0 | 5.52 | 0.917 | 0.989 |
| | | 20% | 10 | 5.0 | 6.44 | 0.804 | 0.969 |
| | | 30% | 15 | 5.0 | 6.78 | 0.767 | 0.961 |
| | 10 | 0% | 0 | 10.0 | 10.00 | 1.000 | 1.000 |
| | | 10% | 5 | 10.0 | 11.16 | 0.901 | 0.972 |
| | | 20% | 10 | 10.0 | 12.68 | 0.798 | 0.935 |
| | | 30% | 15 | 10.0 | 13.84 | 0.736 | 0.906 |
| | 15 | 0% | 0 | 15.0 | 15.00 | 1.000 | 1.000 |
| | | 10% | 5 | 15.0 | 16.70 | 0.900 | 0.953 |
| | | 20% | 10 | 15.0 | 18.60 | 0.812 | 0.900 |
| | | 30% | 15 | 15.0 | 21.10 | 0.719 | 0.831 |
| 100 | 1 | 0% | 0 | 1.0 | 1.00 | 1.000 | 1.000 |
| | | 10% | 10 | 1.0 | 1.14 | 0.930 | 0.999 |
| | | 20% | 20 | 1.0 | 1.22 | 0.903 | 0.998 |
| | | 30% | 30 | 1.0 | 1.20 | 0.907 | 0.998 |
| | 3 | 0% | 0 | 3.0 | 3.00 | 1.000 | 1.000 |
| | | 10% | 10 | 3.0 | 3.36 | 0.912 | 0.996 |
| | | 20% | 20 | 3.0 | 3.48 | 0.884 | 0.995 |
| | | 30% | 30 | 3.0 | 4.04 | 0.807 | 0.989 |
| | 5 | 0% | 0 | 5.0 | 5.00 | 1.000 | 1.000 |
| | | 10% | 10 | 5.0 | 5.64 | 0.900 | 0.993 |
| | | 20% | 20 | 5.0 | 5.84 | 0.873 | 0.991 |
| | | 30% | 30 | 5.0 | 6.78 | 0.771 | 0.981 |
| | 10 | 0% | 0 | 10.0 | 10.00 | 1.000 | 1.000 |
| | | 10% | 10 | 10.0 | 11.00 | 0.915 | 0.989 |
| | | 20% | 20 | 10.0 | 11.98 | 0.849 | 0.978 |
| | | 30% | 30 | 10.0 | 14.52 | 0.709 | 0.950 |
| | 15 | 0% | 0 | 15.0 | 15.00 | 1.000 | 1.000 |
| | | 10% | 10 | 15.0 | 16.52 | 0.913 | 0.982 |
| | | 20% | 20 | 15.0 | 18.22 | 0.829 | 0.963 |
| | | 30% | 30 | 15.0 | 21.70 | 0.706 | 0.922 |

**Table S1**: Sensitivity and specificity analysis results, with a complete set of $N$ and $k$ parameters' values.

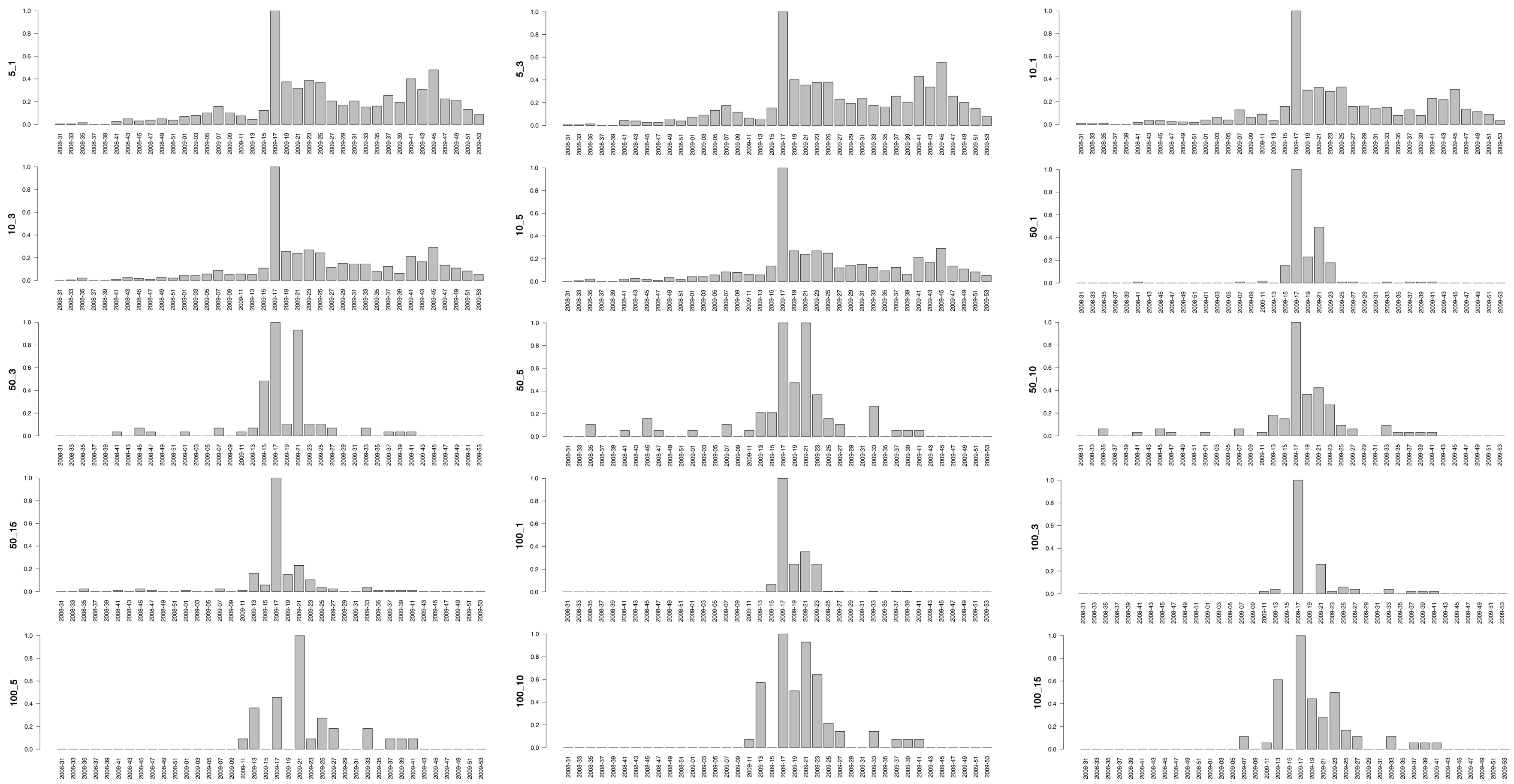


**Figure S1**: Bar plots reporting the warnings for different choices of $N\_k$. Normalized number (over the maximum number) of sequences associated with warnings in H1N1, using different parameters of `stray`. A total of 15 distinct configurations are displayed. In every barplot, intervals of time are on the X-axis, while the maximum value-scaled proportion of sequences labeled as warnings is represented on the Y-axis (min 0, max 1).

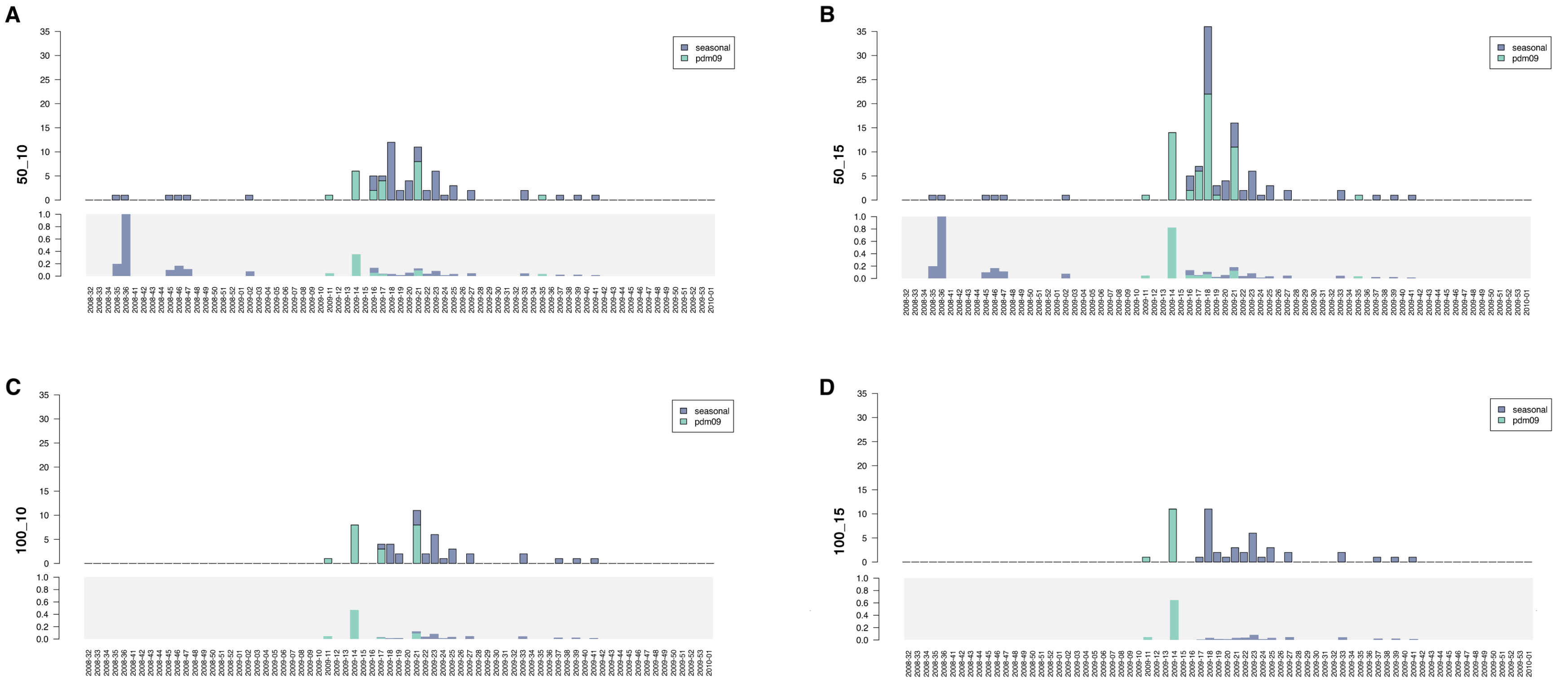


**Figure S2**: Total number of warnings and relative proportion of warnings to the total number of sequences for H1N1 by `stray` with (**A**) $N = 50$ and $k = 10$, (**B**) $N = 50$ and $k = 15$, (**C**) $N = 100$ and $k = 10$, and (**D**) $N = 100$ and $k = 15$. For every configuration, the upper panel illustrates the total number of warnings, while the lower panel indicates the relative proportion w.r.t the total number of sequences. The complete interval of time considered by our analyses is displayed. Colors indicate different viral clades, pdm09 or seasonal.

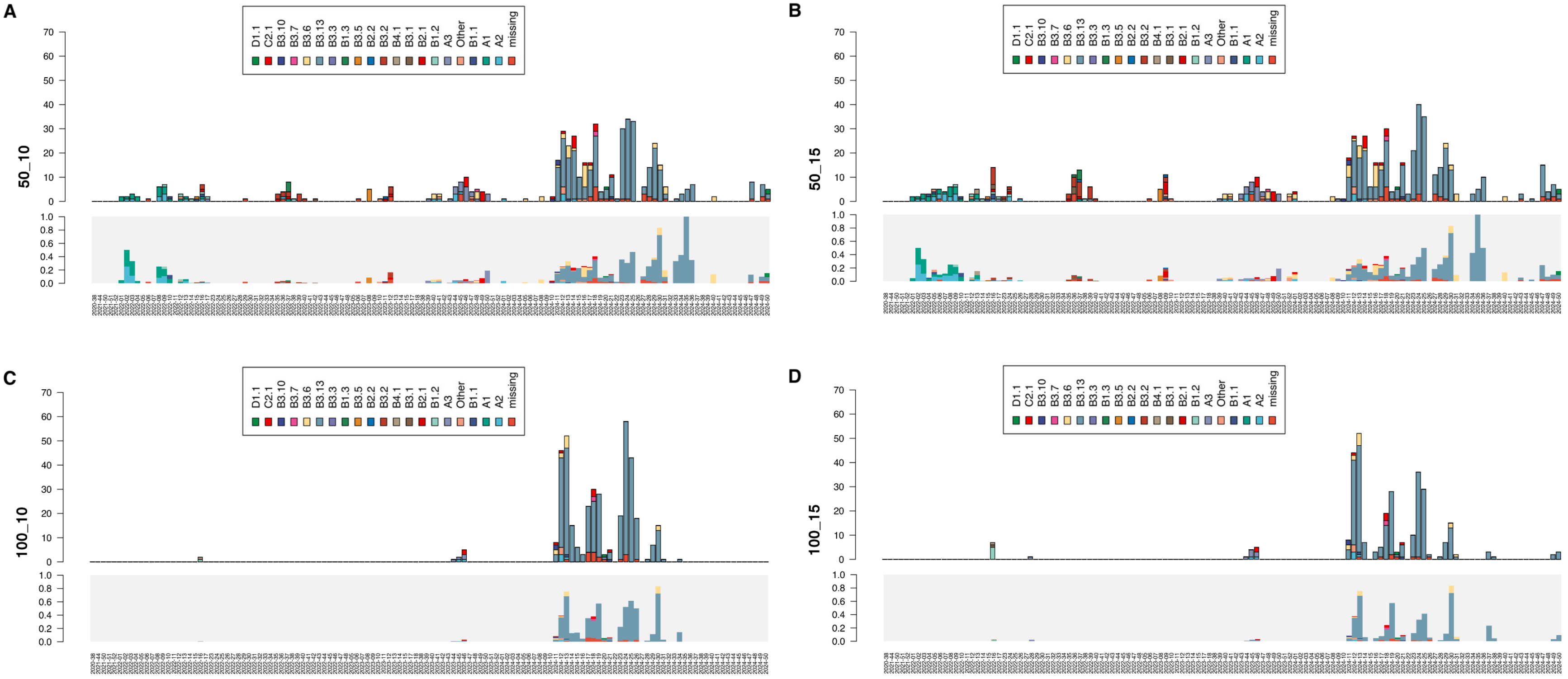


**Figure S3**: Total number of warnings and relative proportion of warnings to the total number of sequences for H5N1 by `stray` with (**A**) $N = 50$ and $k = 10$, (**B**) $N = 50$ and $k = 15$, (**C**) $N = 100$ and $k = 10$, and (**D**) $N = 100$ and $k = 15$. For every configuration, the upper panel illustrates the total number of warnings, while the lower panel indicates the relative proportion w.r.t the total number of sequences. The complete interval of time considered by our analyses is displayed. Colors indicate different viral Genotypes. A clear increase in the number and proportion of warnings associated with the B3.13 genotype is clearly observed starting from Week 2024-11.

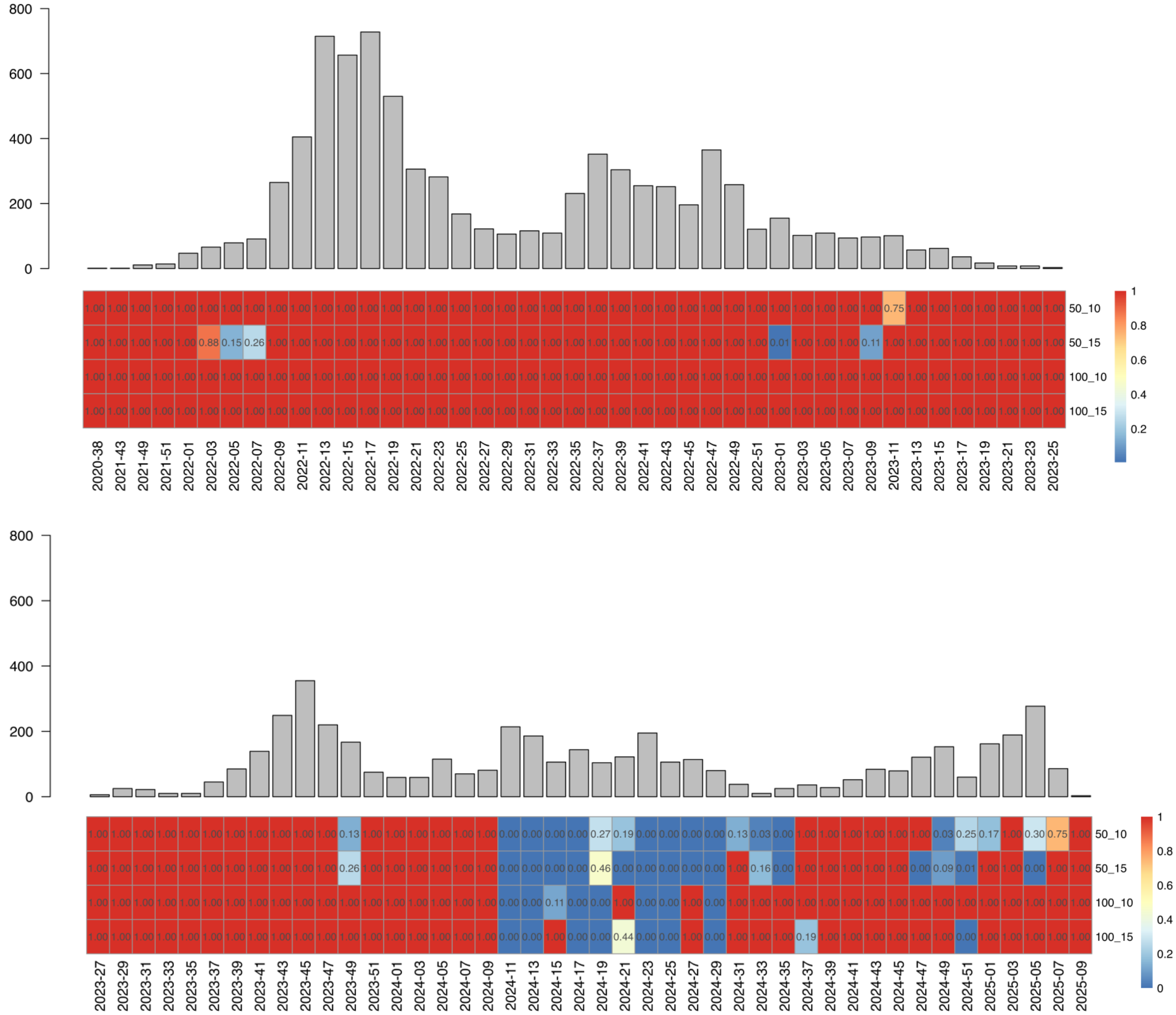


**Figure S4**: *Super Warnings* in H5N1. The heatmap displays FDR-corrected p-values for the statistically significant increase in the number of warnings (Superwarnings) over all the time intervals from 2020-38 to 2025-09. Dark blue cells (corrected p-values ≤ 0.05) indicate *Super Warnings*. Time intervals are shown on columns. Parameters of `stray` on the rows.

**A**

Texas

| Week | 1st day of the Week | N. Sequences | 5 | | 10 | | | 50 | | | | | 100 | | | | |
|---|---|---|---|---|---|---|---|---|---|---|---|---|---|---|---|---|---|
| | | | 1 | 3 | 1 | 3 | 5 | 1 | 3 | 5 | 10 | 15 | 1 | 3 | 5 | 10 | 15 |
| 2024-07 | 2024 Feb 12 | 0 | 0 | 0 | 0 | 0 | 0 | 0 | 0 | 0 | 0 | 0 | 0 | 0 | 0 | 0 | 0 |
| 2024-09 | 2024 Feb 26 | 8 | 5 | 4 | 3 | 2 | 6 | 5 | 0 | 0 | 0 | 0 | 6 | 0 | 0 | 0 | 0 |
| 2024-11 | 2024 Mar 11 | 148 | 53 | 43 | 32 | 42 | 38 | 29 | 38 | 36 | 18 | 14 | 29 | 37 | 31 | 25 | 26 |

Michigan

| Week | 1st day of the Week | N. Sequences | 5 | | 10 | | | 50 | | | | | 100 | | | | |
|---|---|---|---|---|---|---|---|---|---|---|---|---|---|---|---|---|---|
| | | | 1 | 3 | 1 | 3 | 5 | 1 | 3 | 5 | 10 | 15 | 1 | 3 | 5 | 10 | 15 |
| 2024-09 | 2024 Feb 26 | 0 | 0 | 0 | 0 | 0 | 0 | 0 | 0 | 0 | 0 | 0 | 0 | 0 | 0 | 0 | 0 |
| 2024-11 | 2024 Mar 11 | 14 | 3 | 2 | 3 | 5 | 5 | 1 | 4 | 5 | 5 | 5 | 1 | 5 | 7 | 7 | 7 |
| 2024-13 | 2024 Mar 25 | 22 | 5 | 7 | 3 | 4 | 8 | 5 | 5 | 4 | 0 | 0 | 5 | 6 | 5 | 0 | 0 |

Idaho

| Week | 1st day of the Week | N. Sequences | 5 | | 10 | | | 50 | | | | | 100 | | | | |
|---|---|---|---|---|---|---|---|---|---|---|---|---|---|---|---|---|---|
| | | | 1 | 3 | 1 | 3 | 5 | 1 | 3 | 5 | 10 | 15 | 1 | 3 | 5 | 10 | 15 |
| 2024-11 | 2024 Mar 11 | 0 | 0 | 0 | 0 | 0 | 0 | 0 | 0 | 0 | 0 | 0 | 0 | 0 | 0 | 0 | 0 |
| 2024-13 | 2024 Mar 25 | 12 | 2 | 2 | 1 | 1 | 2 | 3 | 5 | 5 | 5 | 5 | 7 | 11 | 11 | 11 | 11 |
| 2024-15 | 2024 Apr 08 | 27 | 3 | 3 | 3 | 2 | 2 | 2 | 3 | 2 | 0 | 0 | 3 | 4 | 4 | 0 | 0 |

Colorado

| Week | 1st day of the Week | N. Sequences | 5 | | 10 | | | 50 | | | | | 100 | | | | |
|---|---|---|---|---|---|---|---|---|---|---|---|---|---|---|---|---|---|
| | | | 1 | 3 | 1 | 3 | 5 | 1 | 3 | 5 | 10 | 15 | 1 | 3 | 5 | 10 | 15 |
| 2024-13 | 2024 Mar 25 | 0 | 0 | 0 | 0 | 0 | 0 | 0 | 0 | 0 | 0 | 0 | 0 | 0 | 0 | 0 | 0 |
| 2024-15 | 2024 Apr 08 | 5 | 3 | 2 | 2 | 4 | 5 | 3 | 4 | 5 | 0 | 0 | 4 | 4 | 5 | 0 | 0 |
| 2024-17 | 2024 Apr 22 | 6 | 3 | 3 | 3 | 3 | 5 | 6 | 6 | 6 | 6 | 6 | 3 | 5 | 6 | 1 | 0 |

**B**

California

| Week | 1st day of the Week | N. Sequences | 5 | | 10 | | | 50 | | | | | 100 | | | | |
|---|---|---|---|---|---|---|---|---|---|---|---|---|---|---|---|---|---|
| | | | 1 | 3 | 1 | 3 | 5 | 1 | 3 | 5 | 10 | 15 | 1 | 3 | 5 | 10 | 15 |
| 2024-33 | 2024 Aug 12 | 0 | 0 | 0 | 0 | 0 | 0 | 0 | 0 | 0 | 0 | 0 | 0 | 0 | 0 | 0 | 0 |
| 2024-35 | 2024 Aug 26 | 21 | 3 | 5 | 3 | 6 | 6 | 3 | 7 | 0 | 8 | 11 | 3 | 7 | 9 | 0 | 0 |
| 2024-37 | 2024 Sep 09 | 31 | 15 | 7 | 13 | 10 | 9 | 14 | 9 | 4 | 0 | 0 | 15 | 24 | 18 | 0 | 3 |
| 2024-39 | 2024 Sep 23 | 20 | 8 | 8 | 10 | 3 | 3 | 7 | 6 | 0 | 0 | 0 | 7 | 7 | 0 | 0 | 0 |
| 2024-41 | 2024 Oct 07 | 21 | 15 | 10 | 11 | 10 | 8 | 10 | 7 | 0 | 0 | 0 | 18 | 13 | 0 | 0 | 0 |
| 2024-43 | 2024 Oct 21 | 19 | 10 | 10 | 9 | 10 | 9 | 12 | 0 | 0 | 0 | 2 | 15 | 0 | 0 | 0 | 0 |
| 2024-45 | 2024 Nov 04 | 22 | 14 | 13 | 12 | 14 | 13 | 7 | 0 | 0 | 0 | 1 | 12 | 0 | 0 | 0 | 0 |
| 2024-47 | 2024 Nov 18 | 46 | 25 | 19 | 20 | 27 | 23 | 9 | 4 | 3 | 8 | 18 | 6 | 0 | 0 | 0 | 0 |
| 2024-49 | 2024 Dec 02 | 14 | 10 | 7 | 8 | 9 | 11 | 6 | 3 | 6 | 7 | 7 | 6 | 0 | 0 | 0 | 5 |
| 2024-51 | 2024 Dec 16 | 11 | 8 | 7 | 8 | 8 | 7 | 9 | 4 | 2 | 5 | 9 | 8 | 0 | 2 | 3 | 7 |
| 2025-01 | 2024 Dec 30 | 27 | 14 | 9 | 11 | 9 | 10 | 16 | 3 | 4 | 4 | 4 | 18 | 2 | 2 | 2 | 3 |
| 2025-03 | 2025 Jan 13 | 17 | 15 | 12 | 10 | 10 | 10 | 0 | 0 | 2 | 3 | 3 | 1 | 0 | 1 | 3 | 3 |
| 2025-05 | 2025 Jan 27 | 31 | 13 | 18 | 14 | 20 | 25 | 17 | 4 | 1 | 17 | 18 | 20 | 4 | 0 | 3 | 9 |
| 2025-07 | 2025 Feb 10 | 2 | 2 | 2 | 1 | 2 | 2 | 2 | 0 | 2 | 1 | 1 | 2 | 0 | 0 | 1 | 1 |
| 2025-09 | 2025 Feb 24 | 0 | 0 | 0 | 0 | 0 | 0 | 0 | 0 | 0 | 0 | 0 | 0 | 0 | 0 | 0 | 0 |
| 2025-11 | 2025 Mar 10 | 0 | 0 | 0 | 0 | 0 | 0 | 0 | 0 | 0 | 0 | 0 | 0 | 0 | 0 | 0 | 0 |
| 2025-13 | 2025 Mar 24 | 0 | 0 | 0 | 0 | 0 | 0 | 0 | 0 | 0 | 0 | 0 | 0 | 0 | 0 | 0 | 0 |
| 2025-15 | 2025 Apr 07 | 0 | 0 | 0 | 0 | 0 | 0 | 0 | 0 | 0 | 0 | 0 | 0 | 0 | 0 | 0 | 0 |
| 2025-17 | 2025 Apr 21 | 0 | 0 | 0 | 0 | 0 | 0 | 0 | 0 | 0 | 0 | 0 | 0 | 0 | 0 | 0 | 0 |

**Figure S5**: (**A**) Warnings emitted for Texas, Michigan, Idaho, and Colorado, marking the progression of the diffusion of warnings from February 12th, 2024, to April 22nd, 2024. (**B**) Warnings emitted for California from August 12th, 2024, to April 21st, 2025. Every row reports bi-weekly counts of warnings, according to all possible assignments of parameters $N$ and $k$.

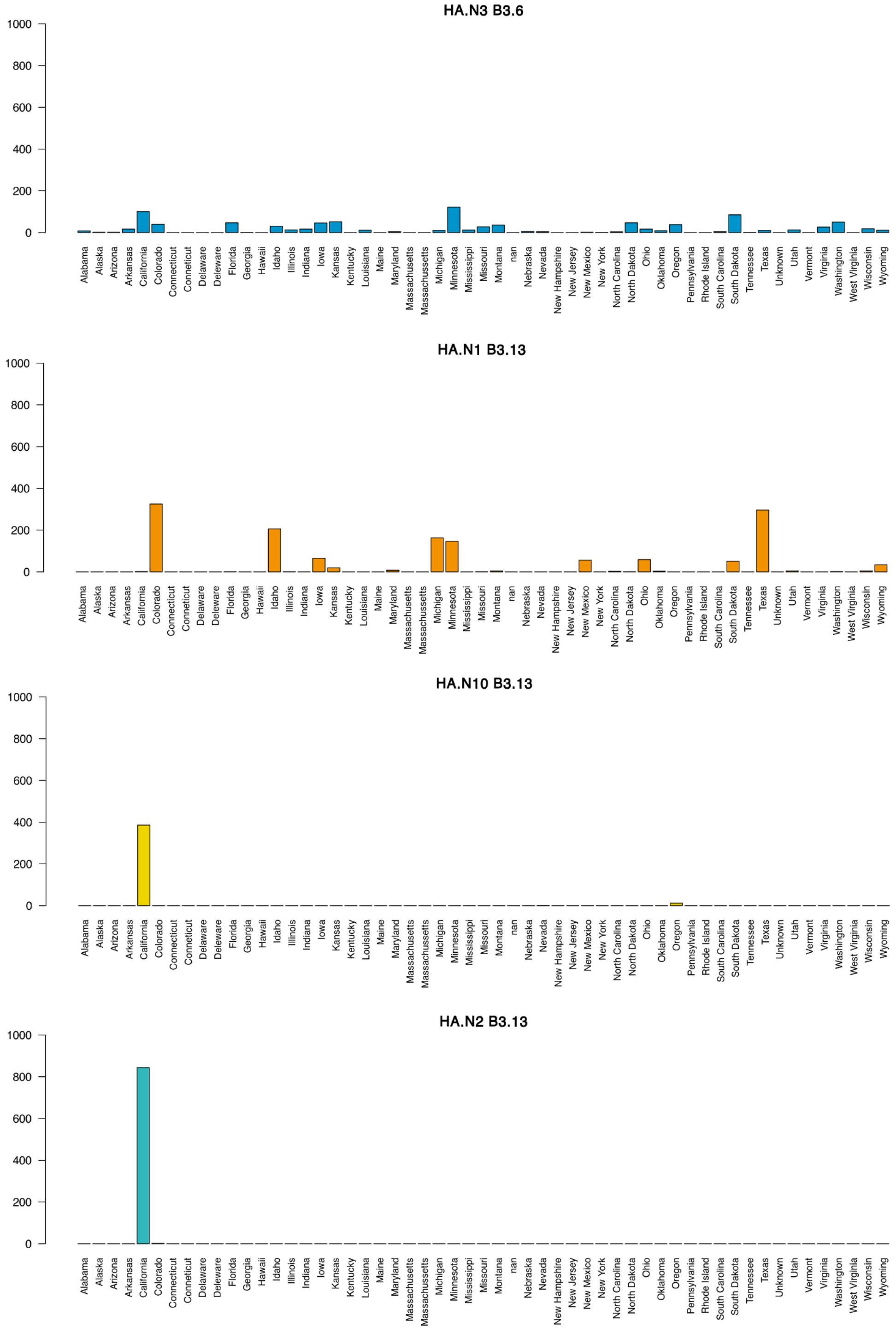


**Figure S6**: Geographic distribution of selected HaploCoV clusters in the United States. The number of sequences is represented on the Y-axis, and States are on the X-axis. Only HaploCoV designations corresponding with the B3.13 (HA.N1, HA.N2, and HA.N10) genotype and its ancestor B3.6 (HA.N3) are displayed. While HA.N3 and HA.N1 display more widespread distributions, HA.N2 and HA.N10 are observed exclusively in dairy cows in California.
